# Demonstration that sublinear dendrites enable linearly non-separable computations

**DOI:** 10.1101/2023.06.01.543223

**Authors:** Romain D. Cazé, Alexandra Tran-Van-Minh, Boris S. Gutkin, David A. DiGregorio

**Author notes:** joint senior authors (Boris S. Gutkin, David A. DiGregorio).

## Abstract

Theory predicts that nonlinear summation of synaptic potentials within dendrites allows neurons to perform linearly non-separable computations (LNSCs). Using Boolean analysis approaches, we predicted that both supralinear and sublinear synaptic summation could allow single neurons to implement a type of LNSC, the feature binding problem (FBP), which does not require inhibition contrary to the XOR. Notably, sublinear dendritic operations enable LNSCs when scattered synaptic activation generates increased somatic spike output. However, experimental demonstrations of scatter-sensitive neuronal computations have not yet been described. Using glutamate uncaging onto cerebellar molecular layer interneurons, we show that scattered synaptic-like activation of dendrites evoked larger compound EPSPs than clustered synaptic activation, generating a higher output spiking probability. Moreover, we also demonstrate that single interneurons can indeed implement the FBP. We use a biophysical model to predict under what conditions a neuron can implement the FBP and what leads to failures. Experimental results agree with the model-determined conditions and hence validate our protocol as a solid benchmark for a neuron to implement linearly non-separable computations. Since sublinear synaptic summation is a property of passive dendrites we expect that many different neuron types can implement LNSCs.

## Introduction

Dendritic filtering influences the shape and amplitude of postsynaptic potentials depending on the synaptic conductance, dendritic morphology, synapse location and the expression of voltage-gated channels^1^. Dendritic properties also influence how multiple synaptic potentials sum, either linearly (if the response amplitude equals the arithmetic sum of the individual synaptic responses), sub-linearly (response amplitude below the sum of the individual responses), or supra-linearly (response amplitude above their sum)^2^. This dendrite-specific arithmetic can greatly enhance a neuron’s computational abilities^3^ and is thought to be prominent in human neurons^4^. Supra-linear summation is thought to underlie active whisker sensation^5^, generate orientation selectivity^6^, grid cell activity^7^, and sensory perception^8^. In contrast, little is known about the computational advantages imparted by sub-linear dendritic integration^9,10^. Recent computational studies suggest that sub-linear integration in cortical fast-spiking interneurons could contribute to memory encoding^11^.

Our previous theoretical work showed a single dendritic non-linearity (either sub- or supralinear) significantly expands the number of functions a neuron can implement^12^.For example, for 8 excitatory uncorrelated inputs, a neuron with a sufficient number of non-linear dendrites (either sub our supralinear) can implement 10^17^ more computations (see methods in^13^ for details). These additional computations are, by definition, a part of the class of linearly non-separable computations (LNSCs). It is interesting to note that passive sublinear dendrites are able to expand the neurons computational capacity on the par with active dendritic nonlinearities, albeit under different synaptic placement conditions (see below).

While LNSCs cannot be implemented by a point neuron model, a neuron with sublinear dendrites can robustly implement an LNSC called the feature binding problem (FBP) (Fig 1). The FBP computation allows a neuron to represent multiple distinct objects, each comprised of independent features. An example of a FBP is given in Figure 1 where the goal is to detect a yellow circle or a green triangle separately by giving an output of 1, but not respond to a yellow triangle or a green circle (Fig 1A). This distinction cannot be done by linearly separating the inputs based on their summed amplitudes. We note that mathematically speaking, aside for the specific feature combinations that must yeild 1 or 0, for the FBP example we consider, there are two feature combinations that do not require specific responses (i.e. two colors or two shapes can give either 0 or 1). Because FBPs can be implemented with only excitatory synapses, we used glutamate uncaging at single synapses to demonstrate this FBP experimentally.

**Figure 1.**
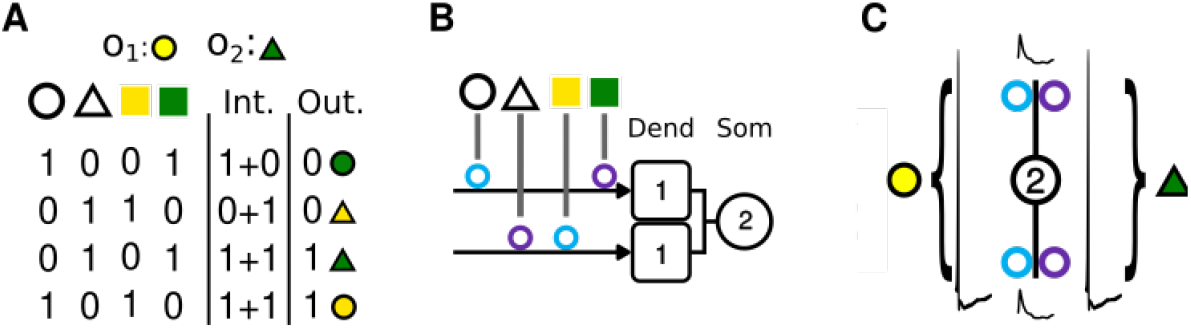
The Feature Binding Problem: A linearly inseparable computation and its implementation in a scatter sensitive neuron. (A) A truth table describing the FBP of two objects with two features each. First column describes presynaptic activity: response of a dendrite is active:1 or not:1. The ntegration column “Int.” describes how a neuron dendrites which implement this computation should integrate inputs. Here we define that the a dendrite saturates at 1. Hence for two clustered inputs the response of the dendrite is 1+0, for scattered it is the arithmetic sum 1+1. The output “Out.” column shows if a neuron spikes:1 or not:0. We omit some inputs because whatever the response to them this computation would still be a LNSC (see Methods for more details). (B) FBP dendritic implementation: both dendrites must be saturated (saturation threshold of 1 in the square) to trigger neuron’s spiking (threshold of 2 in the circle). Because Circle+Green and Triangle+Yellow inputs target the same dendrite, the resulting integration fails to reach the depolarization does not reach the threshold at the soma necessary for the spike. Yet for the two objects the inputs are scattered, both dendrites are saturated and the soma surpasses its threshold. (C) indicates that features activating the same dendrites do not reach threshold and cannot generate a spike, but those features scattered onto the two dendrites generate a somatic spike.

To implement the FBP, supralinear integration requires that the correct feature conjunction for an object cluster on a dendritic section, while feature conjunction that should lead to an absence of response must scatter over the dendritic tree. In contrast, sublinear integration requires the opposite^1,12^: object related conjunctions that should fire the neuron should scattered on different dendritic segments (or dendrites), irrelevant conjuctions should not scatter. Thus the case where scattered inputs fire the cell and clustered inputs do not is essential for solving the FBP using sublinear dendrites (Fig 1C). Here we used glutamate uncaging and biophysical modeling to show that the sublinear dendrites of cerebellar molecular layer interneurons render them scatter-sensitive, thereby enabling them to implement the FBP.

## Results

### Cerebellar stellate cells are scatter sensitive

Dendrites of cerebellar stellate cells are thin and integrate synaptic conductances passively^9^. The high-impedance dendrites generate large local depolarization upon excitatory synaptic activation that reduce the driving force for synaptic currents, resulting in a sublinear summation of postsynaptic potentials when activating multiple synapses^14^. Therefore, when synapses on the same sublinear dendrite are activated simultaneously, they “interact” and thus sum less effectively than if synapses were activated on two different dendrites, due to a lack of effective electrical interaction. In other words, the somatic depolarization is larger and more sensitive to synapse activation patterns that are “scattered” across the dendritic tree.

We used scanning two-photon glutamate uncaging^14,15^ in parasagittal cerebellar brain slices to test the hypothesis that neurons with sublinear dendrites generate larger excitatory potentials when synaptic activation is distributed across dendrites rather than synaptic activation within a dendrite (Fig 2). We identified stellate cells by patching somata located in the outer third of the molecular layer. To mimic clustered activation, we uncaged glutamate using 0.2 ms 720 nm laser pulses in four locations within putative stellate cell dendritic trees in any one trial (slices were superfused wth 2 mM MNI-glutamate) and recorded uncaging-evoked excitatory postsynaptic potentials (uEPSPs) using somatic current clamp patch recordings. Current was injected into the soma to maintain a resting potential near -70mV. Either all uncaging locations were clustered in the same dendrite, or two uncaging locations were placed on two different primary dendrites (see Fig 2A). We used pairs of uncaging sites to increase the local depolarization. Compound subthreshold uEPSPs were generated from four sites ranging from 4-22 mV across all experiments (Fig 2C). We observed that compound uEPSPs evoked on the same dendritic branch were systematically smaller than their linear sum, as expected for sublinear integration (Fig 2B, C). However, compound uEPSPs generated from uncaging locations scattered onto two different dendrites were not significantly different than their linear sum and thus consistent with scatter-sensitive subthreshold PSPs. Because of the inertia of galvanometer mirrors, we could more easily perform simultaneous activation of three dendrites if one was stimulated using an extracellular electrode (see Methods). Results were similar for the activation of effectively six uncaging sites across three dendrites (Fig 2D-F). These results confirm that compound synaptic potentials in neurons with sublinear dendrites were larger if the inputs were scattered across dendrites rather than clustered on a single dendrite.

**Figure 2.**
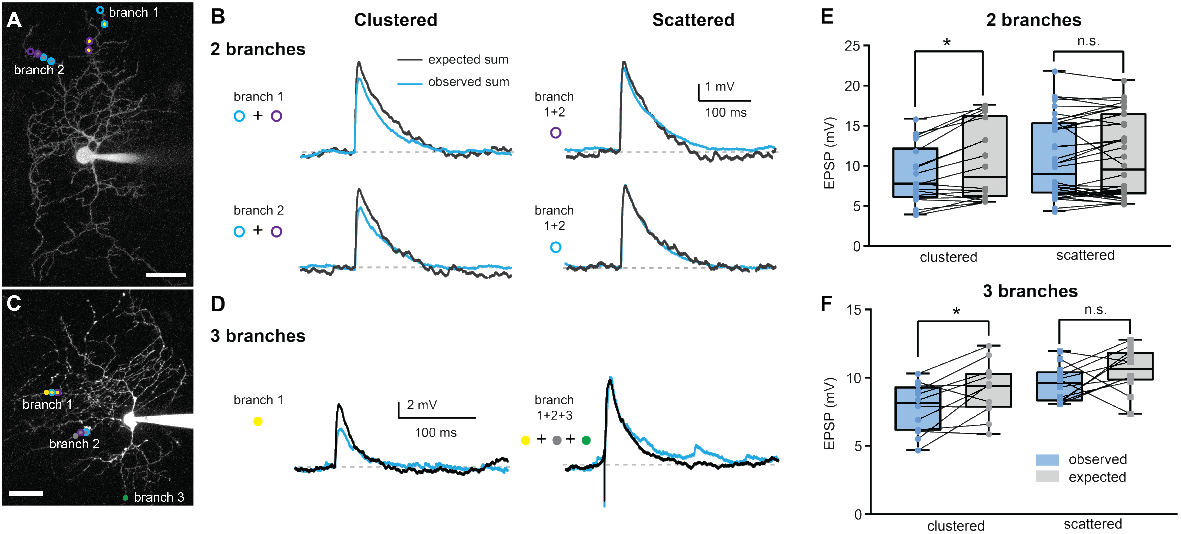
Scattered inputs sum more linearly than clustered inputs. (A, C) 2PLSM image of a cerebellar stellate cell filled with Alexa Fluor 594. Colored spots indicate dendritic locations stimulated using glutamate uncaging (branches 1 and 2, A and C) or electrical stimulation (branch 3, C). (B, D) Example traces of somatic current-clamp recordings in response to stimulation of a group of clustered (left) of scattered (right) inputs (blue traces), and traces obtained as the sum of EPSPs recorded in response to stimulation of individual locations (black traces). (E, F) Peak amplitudes of observed and expected EPSPs in response to stimulation of clustered or scattered inputs.Note that in panel F while the difference between the expected and observed voltage responses are not statistically significant for the number of neurons recorded, we allow the possibility that this may change with a larger number of cells.Note that lines connect responses of a single cell on a single clustered vs scattered trials.

Unlike most studies that focus on the characterization of subthreshold dendritic operations, we set out to verify that the scatter-sensitive subthreshold behavior of stellate cell dendrites also translated into scatter-sensitive spiking probability. We applied the same uncaging protocol described for Fig 2, but adjusted the holding current to maintain the resting potential around -60 mV, to facilitate spiking. Indeed, we found that uncaging locations that were located on the same dendrite produced a significantly lower spiking probability than when uncaging locations were on separate dendrites (Fig 3). These data demonstrate that stellate cells are more likely to fire when inputs are distributed over their dendritic tree than when they cluster on specific dendrites. We also extended this study to three-branch stimulation as described above and observed a robust increased firing probability if the stimulation was distributed across the dendrites (Fig 3C, D, and E). Thus, the neuronal computation (as represented by somatic spiking) of cerebellar stellate cells is scatter-sensitive. It remains to be demonstrated that the FBP can be computed by the neuron with scatter-sensitive spike production.

**Figure 3.**
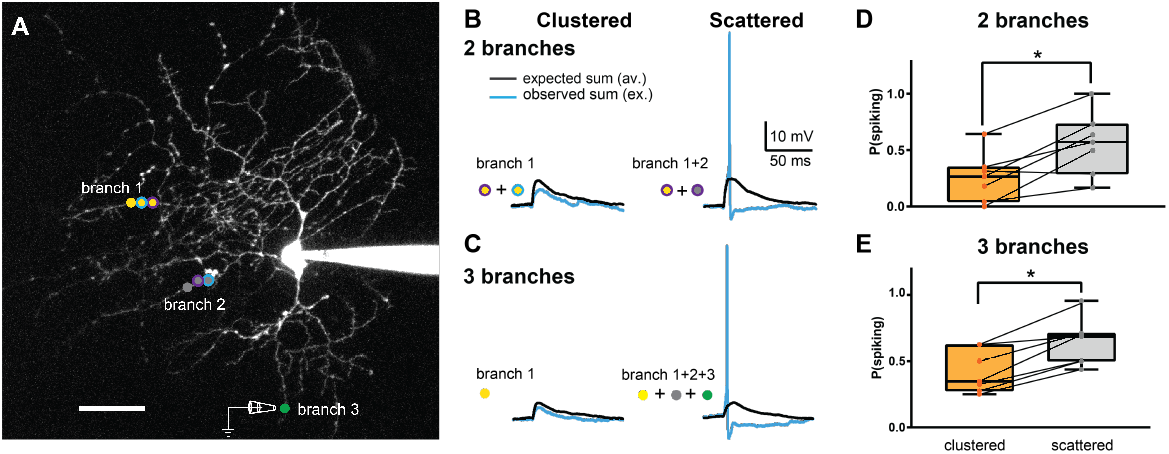
Multi-branch glutamate uncaging reveals a higher firing probability for scattered versus clustered stimulation. (A) 2PLSM image of a cerebellar stellate cell filled with Alexa Fluor 594. Colored spots indicate locations stimulated using glutamate uncaging (branch 1 and 2) or electrical stimulation (branch 3). (B-C) Example traces of somatic current-clamp recordings of EPSPs and action potentials in response to stimulation of a group of clustered (left) or scattered (right) inputs on 2 (B) or 3 (C) branches. (D-E) Spike probabilities in response to clustered:orange and scattered:grey synaptic inputs.

Since we established that stellate cells are scatter-sensitive, we next examined whether such scatter sensitivity would confer them the ability to perform linearly non-separable computations. Specifically, we set out to show they could implement the FBP (see Fig 4). Glutamate was uncaged at four distinct locations in six different patterns. There were four scattered cases (uncaging sites on different dendrites) and two clustered cases (uncaging sites on the same dendrite). To observe an FBP we examined whether a neuron remained silent (no action potential) in the two clustered cases and fired in two scattered cases with non-overlapping uncaging sites. Let us consider each uncaging location as a synaptic input encoding an object’s feature, then combinations of uncaging locations that trigger a somatic spike effectively encode the object. If we observed that the neuron fires for specific combinations of non-overlapping uncaging sites and stays silent for others, then we can claim that it implements the FBP. To give a proof-of-principle demonstration, we chose to label each uncaging location with a unique symbol shape or color: triangle or a circle shape, and green vs. yellow color. Given such labeling, we would conclude that the feature binding problem was correctly implemented if a green triangle and yellow circle were encoded by the soma. In our experiments, we observed multiple trials in which the results were consistent with the FBP, while in others they were not (either scattered synaptic activation did not produce a spike or the clustered case did) (see Fig 4 lower right, red shading). The cell shown in Figure 4 implemented the FBP in nearly half of the trials (n=15), i.e. it fired for two objects with two specific disjoint features (e.g. green triangle or yellow circle) and stayed silent for two other objects made up of other disjoint feature combinations (the green circle and yellow triangle). We also observed, in this cell, trials where a false positive spike was generated (red shading Fig 4). Because of the costly experimental setup we managed to perform suprathreshold experiments only on four cells, where we stimulated a couple of dendrites. Two cells were capable of implementing the FBP 50% of the time. To strengthen this experimental work and explain cases where cells fail to implement the FBP we used a realistic biophysical model.

**Figure 4.**
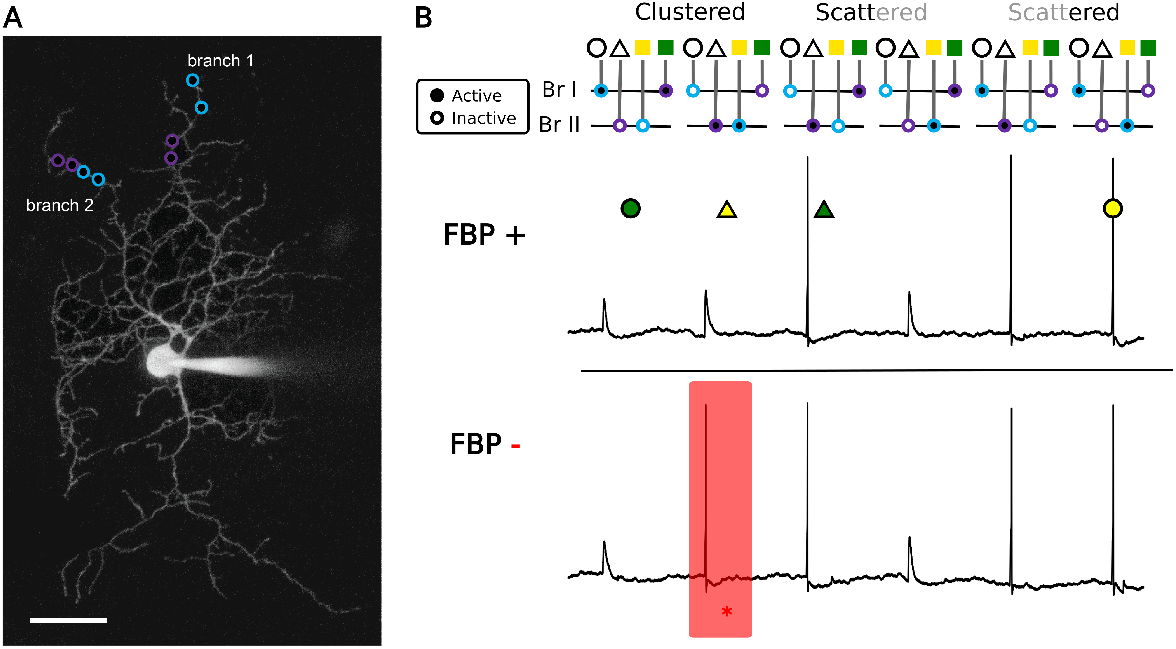
Cerebellar stellate cells might implement the feature binding problem (FBP). (A) 2PLSM image of a cerebellar stellate cell filled with Alexa Fluor 594. Colored spots indicate the uncaging locations, on this cell we uncaged glutamate in four locations, each pair corresponds to one feature (Purple:”Green” on branch 1, “Triangle” on branch 2; Teal:”Yellow” on branch 2, “Circle” on branch 1). The color code is in the top of panel B.and indicates the two objects to be encoded (yellow circle, green triangle). (B) Examples of somatic voltage traces of the same stellate cell in two distinct trials. In the top trial, the neuron computes a feature binding problem, and in the bottom trial it fails to do it because of a false positive (red case). We discarded the two cases when inputs are coming from the shape or the color inputs only as they do not affect the result.

### Understanding why scatter sensitive neurons could compute (or not) the feature binding problem

To understand better the origin of the failed FBP trials, we implemented a biophysical model of a cerebellar stellate cell. We built a multi-compartment model with a realistic dendritic tree morphology (using a reconstruction of cerebellar stellate cell^16^) that was capable of reproducing the uEPSP amplitudes observed in the subthreshold protocol(see Fig 5, 4 pS peak synaptic conductance). The dendrites were modeled as passive and showed sublinear EPSP summation with clustered inputs and more linear summation with inputs scattered on different dendrites (Fig 5A and B). We considered two principal factors that could underlie trial-to-trial variability as well as cell variability in the ability to implement the FBP: EPSP size relative to the threshold and membrane fluctuations. To correctly implement the FBP, the voltage differential from rest to threshold must be such that the single uEPSP and the sublinear compound uEPSP (two-site activation within the same dendrite) are subthreshold, but the linearly summed uEPSPs (two-site activation on different dendrites) are suprathreshold. We thus varied the resting membrane voltage and examined if simulated compound uEPSPs (adjusted to 10 pS per site to ensure spikes) produced false positive or negative spikes (compare Fig 5 to Fig 4). For the depolarized resting membrane potential of -64mV the model reaches the threshold not only in all the scattered stimulation cases but also in one of the clustered cases creating a false positive. When the membrane potential was decreased to -68mV the neuron reached the threshold only in the scattered case, thus correctly implementing the FBP. Finally, if the resting membrane potential was set too low (−72mV), the neuron spiked only in one of the scattered cases. Together, these three simulations exemplify that variations in the resting membrane voltage can influence the successful computation of the FBP. Thus, the EPSP size relative to the threshold must be tuned correctly to implement the FBP.

**Figure 5.**
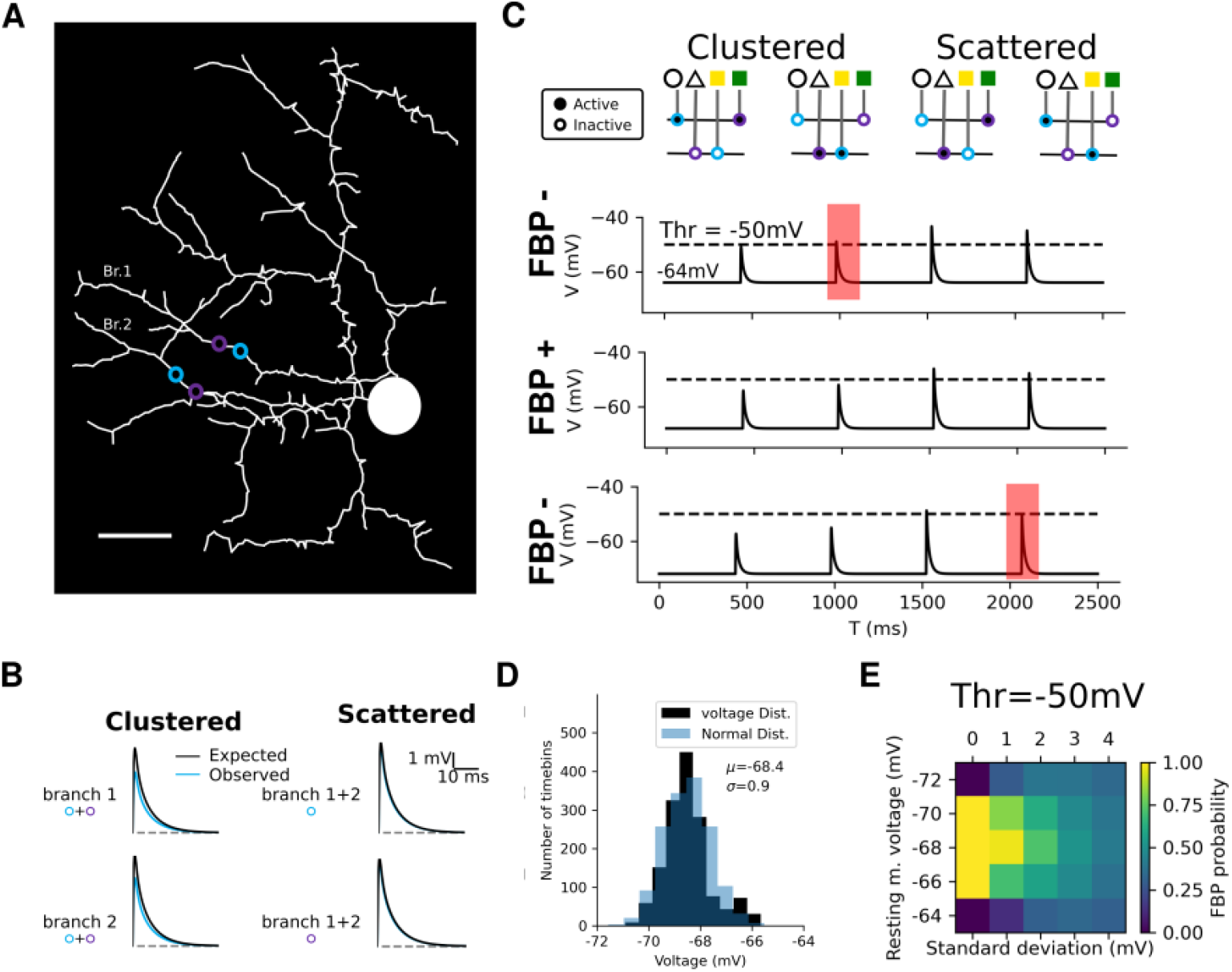
Simulations exploring synaptic and cell parameters necessary to perform FBP. (A) Morphologically detailed model of a cerebellar stellate cell, with colored circles indicating input locations corresponding to the two combinations of features (teal:yellow+circle;purple:green+triangle. (B) compound uEPSPs were simulated by placing two 4 pS synaptic conductance (max value) with a *τ* = 1*ms* time constant on the same (clustered) or different dendrites (scattered). (C) Membrane potential responses for compound uEPSP responses when stimulating pairs of uncaging locations as indicated in the diagram. The dashed line is the the threshold for spike generation. Note that only a resting membrane potential of -68 mV allows for a correct implementation of the FBP, the two other voltages lead to errors in the FBP implementation-being due to respectively a false positive (extraneous spike) and a false negative (absence of a spike). (D) Histogram of the experimental voltage distribution of the voltage (black, we used the first 300 ms without spike and measure the mean voltage in 2 ms timebins) follows a normal like distribution (*μ*=-68.5 mV, *σ* =0.9 mV). We reproduced this distribution using the same mean and standard deviation using a normal distribution (light blue superimposed). (E) Probability of implementing the FBP for different resting membrane potentials and noise levels (SD), calculated from 1000 simulations for each condition.

We also recognized that membrane potential fluctuations could generate false positives and false negatives. We, therefore, modeled the voltage fluctuations using a Gaussian noise distribution that matched experimental observations (mode at -68.4 mV with a standard deviation equal to 0.9 mV vs. resting potential mean for the cell in (Fig 6) was -68.5 mV with an SD of 0.8 mV (Fig 5D)). We, therefore, systematically varied the noise level around this experimental value as well as varying the resting membrane potential of the model from -72mV and -64mV (Fig 5E). To calculate the FBP probability we performed 1000 trials for each simulation condition. As expected from above, when the resting membrane potential is too high the neuron generated false positive spikes, and when it is too low, the neuron generated false negative errors. The simulations indicated an optimum around a resting membrane potential of -68mV, but the FBP probability degraded within increasing membrane potential fluctuations.

**Figure 6.**
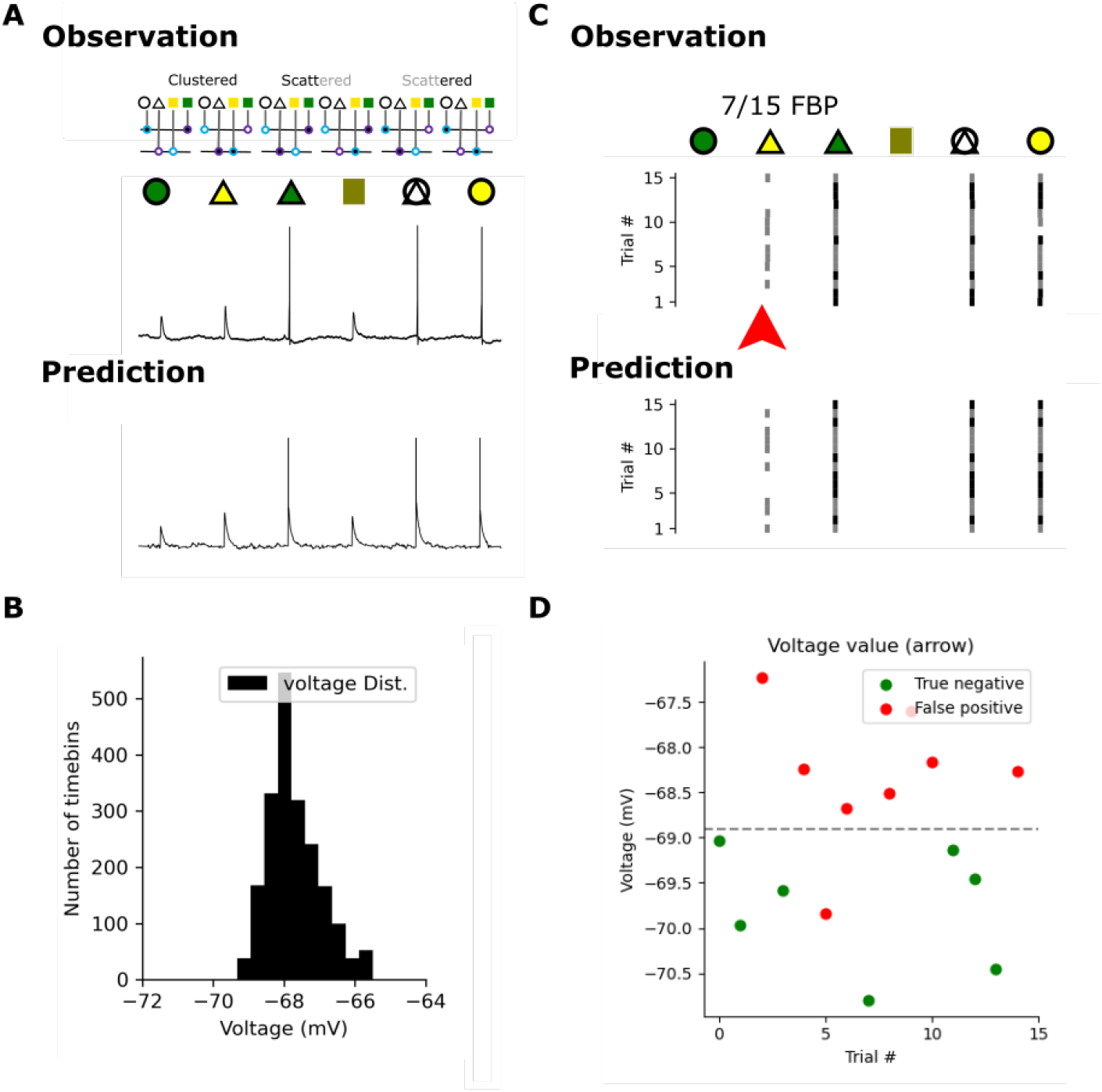
Model simulations reproduce measured FBP probability. (A) Top, somatic voltage trace recorded experimentally of one cell in one trial. Bottom: voltage trace predicted by our model. Importantly, in both cases the neuron implement the FBP (B) Voltage distribution used in our simulation during a 300ms period without stimulation (histogram using 2ms timebins). Note the similarity with panel D from the previous figure. (C) Top, the obtained experimental result from the cell in Fig. 4 with all trials plotted. Trials in black are those in which the FBP was correctly implemented. Bottom, are 15 simulated trials showing a similar FBP probability. (D) Value of the membrane voltage 10ms before the second stimulation episode (red arrow in panel (C)), coloured in green when the neuron stay silent and in red when the neuron spikes

We were able to show that by adjusting the synaptic conductance in the model to match the uEPSP amplitudes for each individual uncaging site, as well as the membrane potential fluctuations, we could reproduce the probability of implementing the FBP (around 50%) as in the cell shown in Fig 4. We, therefore, conclude that scatter-sensitive neurons can perform the FBP, but the relative amplitude of the EPSP versus the voltage-to-threshold and the membrane fluctuations need to be tuned to reliably compute the FBP.

### Benchmarking that a neuron can solve the feature boning problem

Based on the analysis above, we can now demonstrate that a cerebellar stellate cell can indeed perform the FBP even in noisy experimental conditions, as we can see the cell (Fig 6A-C top) can perform the FBP in approximately 50 percent of the trials as is also shown by our model (Fig 6A-C bottom). As we see in our simulations, like the real cell our model also fails to implement the FBP in certain trials because its resting membrane voltage fluctuate randomly (Fig 6B). We note that the model further matches the error trials in the neuron giving spurious spikes at the inputs encoding the yellow triangle (Fig 6C). This enabled us to computationally reproduce the variation observed experimentally (Fig 6C) and observe the neuron implementing the FBP in seven trials over fifteen, thereby matching the experimentally observed variability in FBP implementation. More generally, out of the 13 neurons we subjected to this protocol, we observed two cells that robustly implemented the FBP with a probability of approx 0.5 (cell shown 7/15 trials, cell not shown 6/11 trials).

In Fig 6 we used the same biophysical model as described above with four synaptic conductances placed as indicated in Fig 5A with peak amplitudes of Branch 1 (Br1) teal:7 pS, purple:3 pS; Branch 2 (Br2) teal:20 pS, and purple:10 pS, to match as close as possible the experimental data. The two first stimulation points have a weaker value than the two last as the second uEPSP is larger than the first. The third input with a high value guarantees that the neuron will fire in the third scattered case (20+3pS) while the sum is lower in the second scattered case (10+7pS) keeping the neuron silent in this situation. Thus this model could reproduce the observation (Fig 6A)

We wondered what were the reasons for the much more restricted implementation of the FBP as opposed to a more robust scatter-sensitivity we observed in Fig XX. One reason for failures several of the cells was that the responses at the different stimulation locations were not even: one location gave much larger somatic voltage responses that fired the cell for almost all the trials. Second was that at least one cell did not fire spikes at all and others did not produce enough spikes to access the FBP implementation.

Ideally, the FBP would be implemented in a neuron where every EPSP would be equal for every stimulation point, as in a synaptic democracy scenario^17^. In summary, scatter sensitivity is a necessary, but not sufficient condition for a sublinear dendritic implementation of the FBP.

Furthermore, Fig 6D explains why the neuron fails to perform the FBP only in certain trials. In this panel we plotted the membrane voltage of the cell just before the second stimulation (10ms before), and we colored each point depending on the cell activity (spikes:red, silence:green). We see that that resting potential cases where a neuron gave a false positive spike, the membrane resting potential was too high, and lead to consistent failure to implement the FBP. While for negatives, the resting potential was too low. We also conclude that it is possible to predict when the neuron implement (or not) the FBP. In summary, as we predicted from our modelling analysis above, the resting potential fluctuations combined with membrane noise lead to the FBP implementation failures.

## Discussion

It is well known that dendritic nonlinearities can theoretically increase the computational capacity of neurons because they enable multi-layered information processing. In particular, while there has been fast-growing theoretical literature arguing for universal computing power of dendritic neurons^3,11,12,18^ (bolstered by experimental results on subthreshold dendritic integration^1,2,4,5^ and the potential role of dendritic processes in neuronal tuning properties^8,19–21^), experimental validation of an actual implementation of LNSC by a neuron endowed with nonlinear dendrites, to our knowledge, has not been performed. In this study, we demonstrated using multi-point glutamate uncaging that the sublinear dendritic integration of cerebellar stellate cells leads to larger EPSPs and spiking probabilities when synaptic activation is spread across dendrites (scatter-sensitive) as compared to when the synapses are activated within the same dendritic branch. We also showed that stellate cells can implement the feature binding problem (as previously predicted in^12^). Unlike a LNSC like the XOR, the FBP can be computed using only excitatory synapses. This singular property enabled our demonstration as the XOR would have required a mixture of caged neurotransmitters or a non-monotone neuronal transfer function absent from stellate cell. Moreover, the FBP is of particular interest since neurons implementing it are able to bind multiple feature combinations together into an object^22,23^. Finally, modeling dendritic integration of the thin passive dendrites of cerebellar stellate cells allowed us to identify the biophysical conditions necessary to perform this benchmark.

We have shown here that in order to implement the FBP using excitatory synapses and sublinear dendrites, the two features of a common object must innervate two different dendrites such that their compound EPSP, upon simultaneous activation, is maximized and generates the large spike probability. Otherwise, if two synapses are simultaneously activated within the same dendritic compartment, and their summed local depolarization is large enough to decrease the driving force for ionic currents, the net somatic depolarization will be smaller than the arithmetic sum. Thus clustered synaptic activation produces less depolarization than when the synapses are electronically independent. For a neuron to bind two (or more) features of an object, the features must be distributed across the dendritic tree to maximize the spike probability associated with the object, a computation termed “scatter-sensitive”. However, an important condition is that the scattered activation produces a somatic spike, whereas the equivalent clustered activation must be subthreshold for spike generation. This fine-tuning requires a specific relation between the single synapse EPSP amplitude, the number of synapses associated with the different features, and the difference between the resting membrane potential and the spike threshold. EPSPs that are too strong generate false positives, whereas EPSPs that are too weak generate false negatives spiking representations of objects (Fig 5). Stronger background membrane potential fluctuations will also generate more false positives and negatives on a trial-to-trial basis (see Fig 4 and Fig 6). Synaptic In conclusion, to show that a scatter sensitive neuron can implement a LNSC you need to have strong and almost equal uEPSP and a well controlled resting membrane voltage.

In this study, we emphasized scatter-sensitivity of the neuron as a key factor for the implementation of linearly non-separable computations as it is possible both in active and passive dendrites. Passive thin dendrites have high input impedances and are good candidates for large local depolarizations that generate sublinear summation. Activation of voltage-gated potassium conductances can also generate sublinear integration^10^. Moreover, it is possible that for small numbers of active synapses, their compound EPSP can be large enough to decrease the driving force locally in the dendrite, but still too small to activate nonlinear conductances^24^. In that window of synaptic depolarization, a scatter-sensitive computation can be performed. Moreover, for dendrites that exhibit robust calcium spikes^9^, once the dendritic spike has been generated, additional synaptic depolarization is ineffective in increasing spike probability. Synaptic contacts scattered onto other dendrites must be activated to generate additional somatic depolarization and increased spike probability. Thus both interneurons and principal neurons could exhibit scatter sensitivity and thus use sublinear summation to implement a LNSC^11^, albeit under specific regimes of synaptic and intrinsic cell membrane parameters. Given that scatter sensitivity responses on passive cable properties of dendritic trees, and is not catastrophically deteriorated by active dendritic processes (albeit under different synaptic and intrinsic regimes), it might be a widely generic property of information processing in wide classes of neurons.

## Methods

### Animals and slice preparation

Animal experiments were performed in accordance with the guidelines of Institut Pasteur, France, and all protocols were approved by the Ethics Committee 89 of Institut Pasteur (CETEA; approval DHA180006). Results of this study are reported in accordance with ARRIVE guidelines (https://arriveguidelines.org).

Cerebellar acute slices were prepared from CB6F1 mice (F1 cross of BalbC and C57Bl/6J) of postnatal day P60 to 90. The mice were killed by rapid decapitation (no anesthetic was used), after which the brains were quickly removed and placed in an ice-cold solution containing (in mM): 2.5 KCl, 0.5 CaCl2, 4 MgCl2, 1.25 NaH2PO4, 24 NaHCO3, 25 glucose, 230 sucrose, and 0.5 ascorbic acid bubbled with 95% O2 and 5% CO2. Parasagittal slices (20 0*μ*m thick) were prepared from the dissected cerebellar vermis using a Leica VT1200S vibratome. After preparation, the slices were incubated at 32 degree Celsius for 30 minutes in the following solution (in mM): 85 NaCl, 2.5 KCl, 0.5 CaCl2, 4 MgCl2, 1.25 NaH2PO4, 24 NaHCO3, 25 glucose, 75 sucrose and 0.5 ascorbic acid. Slices were then transferred to an external recording solution containing (in mM): 125 NaCl, 2.5 KCl, 2 CaCl2, 1 MgCl2, 1.25 NaH2PO4, 25 NaHCO3, 25 glucose and 0.5 ascorbic acid, bubbled with 95% O2 and 5% CO2, and maintained at room temperature for up to 7 hours.

### Slice electrophysiology and imaging

Whole-cell current-clamp recordings were performed from stellate cells (SCs) (33^*°*^C – 36^*°*^C) located in the outer third of the molecular layer, using a Multiclamp 700B amplifier (Molecular Devices), and fire-polished thick-walled glass patch-electrodes (tip resistances of 4-6 MΩ). The pipettes were backfilled with an internal solution containing (in mM): 110 K-MeSO3, 40 HEPES, 1 EGTA, 4.5 MgCl2, 0.49 CaCl2, 10 Na-pyruvic acid, 0.3 NaGTP, 4 NaATP, 10 Tris phosphocreatine and 0.04 Alexa Fluor 594 and adjusted to 305 mOsm and pH 7.3. Synaptic responses were filtered at 10 kHz, and digitized at 100 kHz using an analogue-to-digital converter (model NI USB 6259, National Instruments, USA) and acquired with NClamp (www.neuromatic.thinkrandom.com), running in the Igor Pro environment (Wavemetrics). Current was injected to maintain the membrane potential between -70mV and -90mV (after correcting for liquid junction potentials, calculated to be -7 mV using JPCalcW (Barry, 1994; J. Neurosci. Method., 51: 107-116)), and series resistance was compensated by balancing the bridge and compensating pipette capacitance.

Unless otherwise stated, the external solution included 10 *μ*M SR-95531 to block GABAA receptors to avoid confounding results due to partial blockade of GABAA receptors by MNI glutamate, and 50 *μ*M D-AP5 to avoid stimulation of extra-synaptic NMDA receptors.

A pulsed Ti:Sapphire laser (DeepSee, Spectra-Physics) beam tuned at 810 nm was scanned on the preparation using an Ultima microscope (Bruker Fluorescence Microscopy) mounted on an Olympus BX61WI microscope and equipped with a 60x (1.1 NA) water-immersion objective. Simultaneous two-photon fluorescence and Dodt contrast imaging (Luigs and Neumann, Germany) were used to position extracellular stimulating electrodes and uncaging points along spatially isolated dendrites of Alexa Fluor-594-filled SCs, using a transmitted light PMT mounted after the Dodt tube to acquire a laser-illuminated contrast image simultaneously with the 2PLSM image. Alexa Fluor 594 fluorescence was filtered using 640/100 nm bandpass filters (Chroma) and detected using side-on multi-alkali PMTs (3896, Hamamatsu Photonics). In addition to the light collected through the objective, the transmitted infrared light was collected through a 1.4 NA oil-immersion condenser (Olympus), and reflected on a set of substage photomultiplier tubes (PMTs).

### Chemicals

D-AP5 (D-(−)-2-Amino-5-phosphonopentanoic acid) and SR 95531 (2-(3-Carboxypropyl)-3-amino-6-(4 methoxyphenyl) pyridazinium bromide) were purchased from Abcam, UK. MNI-glutamate (4 methoxy-7-nitroindolinyl caged L-glutamate) was purchased from Tocris Bioscience, UK. Alexa Fluor 594 was from Life Technologies, USA. All other chemicals were purchased from Sigma-Aldrich, France.

### Glutamate uncaging

MNI-glutamate was bath applied at a final concentration of 2 mM in ACSF, and recycled. The solution was kept protected from light and any lamps used for ambient light and microscope trans-illumination used during approach of the patch-clamp electrode were covered with a UV yellow filter to prevent undesired photolysis of MNI-glutamate. The preparation was illuminated through a second set of galvanometer-based scan mirrors, allowing independent and rapid positioning of the photolysis beam. The photolysis laser was either a 1P laser at 405 nm diode laser (Omicron) or a pulsed Ti:Sapphire laser (DeepSee, Spectra-Physics) tuned at 720nm. The outputs of the two lasers were independently modulated to combine uncaging of MNI-glutamate and imaging morphology. The imaging laser beam was modulated using a Pockels cell (350-50-BK 02, Conoptics, Danbury, CT). For 2P uncaging, the intensity and duration (200-300 *μ*s) of the photolysis pulse was modulated using an acousto-optic modulator (MT110-B50A1.5-IR-Hk, AA Opto-Electronic, France). For 1P uncaging, the intensity and duration of the photolysis pulse were. A telescope placed on the path of each uncaging beam (Thorlabs) was used to adjust the convergence angle to both backfill the objective and match the focal plane of the two-photon excitation for imaging. Parfocality of the two beams was verified using bleached spots on a microscope slide coated with fluorescent ink. Photolysis laser powers, estimated at the exit of the objective were <1 mW for 1P uncaging and <20 mW for 2P uncaging. We uncaged near simultaneously at up to 6 spots, each 3-6 *μ*m apart, by rapidly switching the focal spot to each new location along the dendrite at 200 *μ*s intervals. For multi-dendrites stimulation, the minimum displacement time of uncaging mirrors between two uncaging locations constrained the distance between the stimulated dendrites (typically in the order of 100-150 *μ*m between the most distant uncaging locations).

### Distribution of the resting membrane voltage

The voltage membrane is sampled at 1kHz as we would we be incapable to obseve spiking at a lower sample rate. We looked at all the 15 trials and used the first 100ms to plot an histogram of the voltage distribution. We have overlapped surrogate data (n=150000 points) generated from a normal distribution using the observed *μ*=68.48 and *σ* =0.90. We have used a visual verification to test the normality of the distribution as usual statistical test (Shapiro wilk or Kolmogorov) were not adapted for the extended number of data points and as we approximated the distribution as a normal.

### Formal definition of the FBP

The definition of the FBP in Tab.1 ensures that FBP is positive (thefore monotone) and linearly non-separable. In another article, we prove using a constructive proof that that a neuron with a sufficient number of sub or supra-linear subunit could compute all positive functions^13^. While for eight variables there are 2730166 threshold positive computations^25^ the total number of positive computations is non (the 8th dedekind number). This gives the ratio of 10^16^

**Table 1.** The FBP truth table organised depending on the number of 1s in the input vectors (the In row). For input vectors with less than two 1s the neuron should stay silent, and if it has more than two 1s the neuron should fire. When there are two 1s, the neuron should fire for two disjoint conjunctions and stay silent for two of other disjoint conjunctions. This guarantees that the FBP is a LNSC. In the two other cases the neuron could fire or not (meaning of *?*)

| In | 0000 | 0001 | 0010 | 0100 | 1000 | 1001 | 0110 | 0101 | 1010 | 0011 | 1100 | 1110 | 1101 | 1011 | 0111 | 1111 |
| --- | --- | --- | --- | --- | --- | --- | --- | --- | --- | --- | --- | --- | --- | --- | --- | --- |
| Out | 0 | 0 | 0 | 0 | 0 | 1 | 1 | 0 | 0 | ? | ? | 1 | 1 | 1 | 1 | 1 |

It is important to note here that the FBP is a monotone function, contrary to the XOR, therefore it enables our demonstration without using a mixture of excitatory and inhibitory caged neurotransmitter, as cerebellar stellate dendrites behave in a monotone way.

We may have wanted to study a more “complex “ FBP with more features or objects. However, the number of dendrites necessary to compute them could have grown rapidly. We studied this scaling in^13^, and here we focus on the simplest FBP (2 objects and 4 features). This choice is motivated by experimental constraints.

For instance, we could have defined a FBP with four features per object. Let us imagine that such feature dimensions are: size, color, texture, shape. For such FBP, we need to define four cases: two where the neuron fires for a small-green-stripy-triangle (object 1) or for a large-yellow-dotted-circle (object 2) and two where the neuron should stay silent, e.g. for a large-green-stripy-triangle or a small-yellow-dotted-circle. In this case, you need a neuron with a minimum of four dendrites to scatter the features of each object on them, and to prevent spiking you need then to cluster the sizes and another set of features on two different branches (like big and green on one branch and small and yellow on another). Note that such clustering does not need to be exquisitely precise, but sufficient such that these non-spiking conjunctions would saturate the dendritic response and not cause a spike. Since our goal in this manuscript was to provide an experimental demonstration that a dendritic neuron can implement the FBP, we would need to set up an experiment with 4 stimulated dendrites with 4 uncaging locations each. This is beyond experimental capabilities. Hence we focused on the 2-feature-object case.

### Biophysical model

All simulations were performed with Brian 2^26^ and we used a detailed reconstruction from^16^. The code needed to reproduce the last figure is available under an open-source licence. We used a conductance-based model with the reconstructed dendritic arbor. The axial resistance between compartments equals 150Ω. Every compartment contained the following passive currents:

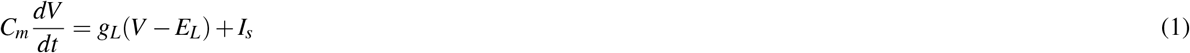

*V* is the membrane potentials, *C*_*m*_ = 1 μFcm^*−*2^ is the membrane capacitance, *g*_*L*_ is the leak capacitance equal to 5*e−* 5siemens which is equivalent to a *R*_*i*_ = 20000Ω input resistance.

The synaptic current *I*_*s*_ is described by

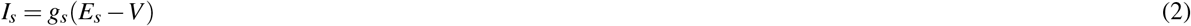

with *E*_*s*_ being the synaptic reversal potential and *g*_*s*_ the synaptic conductance. This conductance jumps up instantaneously for each incoming input and decays exponentially with time constant *τ*_*s*_ = 1 ms otherwise:

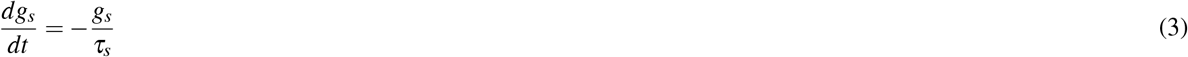

The neuron “spikes” when it reaches -50mV from below and we signal a spike using a vertical bar. We set the resting membrane voltage to -68mV.

### Simulations to match the experimental observation

In Fig 6 we used the same biophysical model as described above with four synaptic conductances placed as indicated (Fig 6A) with peak amplitudes of Br1 teal:7 pS, purple:3 pS; Br2 teal:20 pS, and purple:10 pS exponentially decaying with a 1 ms time constant, to match as close as possible the experimental data (uEPSP amplitude or spike probability). The two first stimulation points have a weaker value than the two last as the second uEPSP is larger than the first. The third input with a high value guarantees that the neuron will fire in the third scattered case (20+3pS) while the sum is lower in the second scattered case (10+7pS) keeping the neuron silent in this situation.

To mimic the observed variation in the membrane resting voltage we used two conductances based processes, one inhibitory and one excitatory of 0.2pS targeting the soma. To create randomly distributed membrane voltage in the model we used the following procedure. We randomly picked 400 integers between 0 and 2000 each with the same probability. Of these 200 corresponded to the times at which an excitatory synapse activates (with the reversal potential of 0mV) and the other 200 to the activation times of an inhibitory synapse (reversal potential -140mV). Both of these surrogate synapses target the soma and have a conductance strengths of 0.2nS. Each synapse was modeled as an instantaneous increase preceding an exponential decay with a 1ms time constant. The goal was to obtain a resting potential distribution that matched the experimentally observed one (see Fig 6A).

The python code to entirely reproduce the two last figures and subpanels is available on zenodo^27^.

## Acknowledgements

B.S.G. would like to acknowledge the support from the Basic Science Program Basic Research Program at the National Research University Higher School of Economics (HSE University). This work was partially supported by the ANR projects “CerebComp” and “InTempCode”.

## Author contributions statement

R.C., A.T. and D.A.D. conceived the experiments, R.C., D.A.D. and B.S.G. conceived the theory, R.C. carried out the computational research A.T. conducted the experiments, R.C. and A.T. analysed the results, D.A.D, B.S.G and R.C. wrote the manuscript. All authors reviewed the manuscript.

## Data availability statement

The datasets used and/or analysed during the current study available from the corresponding author on reasonable request.

## Additional information

To test experimentally our prediction we needed to maximise the neuron’s scatter sensitivity. We used a ball and stick model with a spherical soma (10*μ*m) and two dendrites (200*μ*m length) and then varied the two inputs’ locations (between 50 and 200*μ*m) and the dendritic diameters (between 0.4 and 1*μ*m). Each time we subtracted the result of the scatter (on two dendrites) and of the cluster (on one) stimulation to obtain Fig 7A. This shows how the difference in somatic voltage evolves with the morphological properties of a neuron.

**Figure 7.**
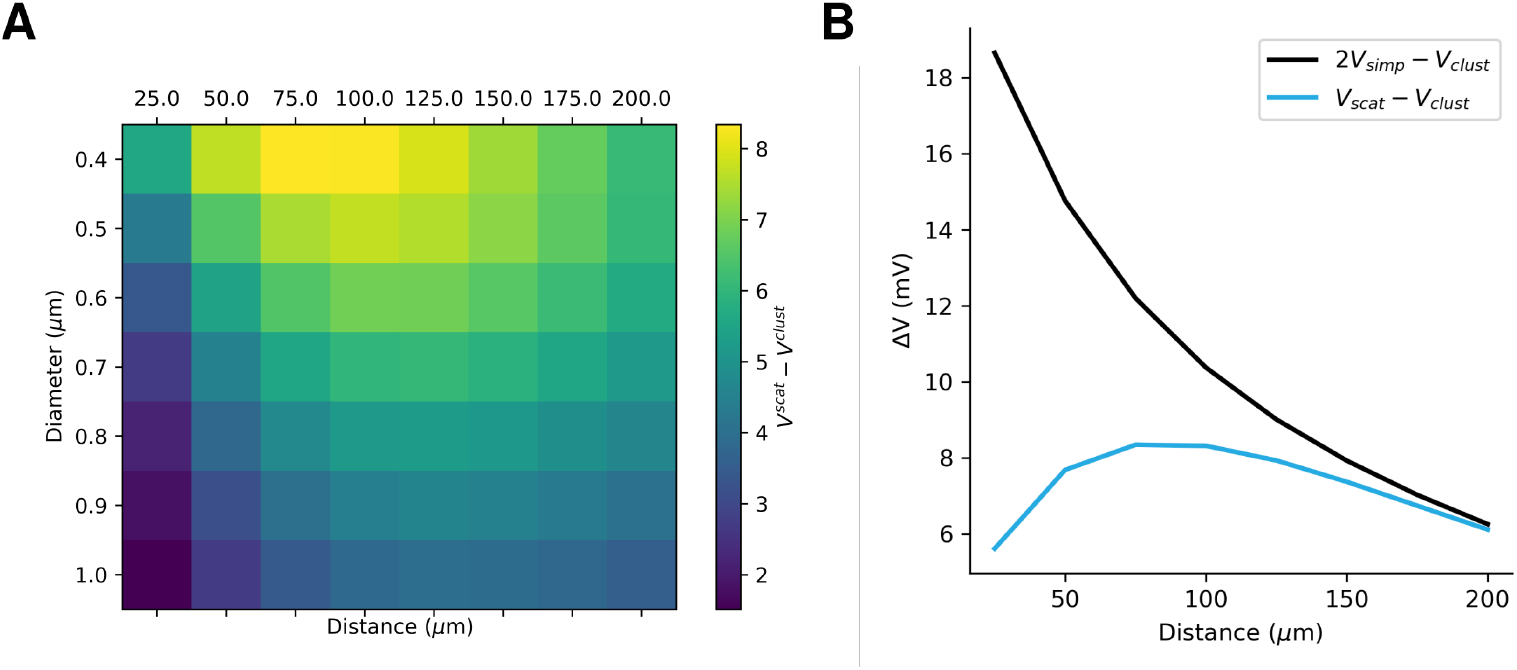
The optimal stimulation location to test our experimental prediction. (A) The difference in the recorded somatic voltage for clustered and scattered synaptic input. Note the optimal at 75 *μ*m.(B) The evolution of the voltage difference for a fixed diameter (0.4*μ*m) in blue and in black the difference between two times the EPSP of a single input and scattered inputs. Note that for a distance superior to a 100 microns the two curves superimpose because of the filtering effect.

One can observe on Fig 7A that the difference is the largest at 75 *μ*m. This optimum depends on the morphological and biophysical parameters of the model. Yet it always exists whatever the parameter set. Still one might wonder the reason for this optimum. Fig 7B explains it. Without surprise the further we were the larger was the difference (as close-by synaptic inputs interact via the soma). Surprisingly we found that after a certain distance this difference decreases. This is because dendritic filtering decrease the effect of both clustered and scattered stimulation on the soma. It leads to a decreased difference between the two types of stimulation.

